# 3D structure of fibroblasts and macrophages in the healthy and cryo-ablated heart

**DOI:** 10.1101/2023.11.30.569388

**Authors:** Marbely C Fernández, Eike M Wülfers, Josef Madl, Stefanie Perez Feliz, Peter Kohl, Callum M Zgierski-Johnston, Franziska Schneider-Warme

**Affiliations:** Institute for Experimental Cardiovascular Medicine - University of Freiburg and Faculty of Medicine, University of Freiburg, Freiburg, Germany

**Keywords:** fibroblasts, macrophages, tissue clearing, 3D reconstruction, cardiac remodelling

## Abstract

**Introduction:** Cardiac non-myocytes (NM) play important roles in heart development, homeostasis, and lesion repair. To assess the relevance of different NM populations for cardiac (patho)physiology, a quantitative assessment of their abundance and structure in the different heart chambers is an essential prerequisite. We here present an experimental approach to determine the distribution, dimensions, and 3D morphology of fibroblasts (FB) and macrophages (MΦ) in healthy and pathologically remodelled hearts.

**Methods and results:** We used Cre-*loxP* recombination to selectively target channelrhopsin-2 (ChR2)-eYFP to either FB or MΦ in healthy and cryo-ablated mouse hearts. Hearts were optically cleared using X-CLARITY and membrane-bound eYFP fluorescence was recorded by confocal microscopy. The resulting image stacks were segmented to generate 3D reconstructions of labelled cell populations in *near native tissue*. In doing so, we show that FB and MΦ have similar surface areas, volumes and morphologies, but that FB occupy larger fractional volumes than MΦ in all chambers of healthy murine hearts. Furthermore, MΦ appear primarily as single cells, whereas FB form extended networks of interconnected cells. In left-ventricular tissue following cryo-ablation, we observed large disordered networks of FB in the scar area with an increased volume occupied by FB both in the scar and remotely. In cryo-ablated ventricles, MΦ form comparatively small, but dense networks in the scar without changing their abundance in remote myocardium.

**Conclusions:** Our study assesses the 3D distribution and structure of fluorescently labelled FB and MΦ in healthy and lesioned murine hearts. Based on 3D reconstructions of FB and MΦ networks, we quantified the surface areas and volumes of individual non-myocytes in the different chambers of the heart and in ventricular scar tissue, thus providing important quantitative data serving as basis for computational modelling of non-myocyte contributions to cardiac structure and physiology.

## 1. Introduction

The heart is a heterocellular organ composed of cardiomyocytes (CM) and non-myocytes (NM). The latter include fibroblasts (FB), endothelial, perivascular and immune cells, and cells of the intracardiac nervous system. CM generally occupy more than half of the myocardial volume and drive the electrical and mechanical activity of the heart. However, while NM occupy a comparatively smaller volume fraction, they account for the majority of the cells. NM play essential roles in cardiac development, maintenance of myocardial structure and function, and cell-cell signalling (Perbellini *et al*., 2018; Lother & Kohl, 2023). Following myocardial injury, NM are activated, proliferate, and invade the scar region from both intra- and extracardiac sources, contributing to scar formation and re-establishment of tissue homeostasis. FB and MΦ represent key NM cell types driving and regulating cardiac lesion repair. In response to injury, cardiac MΦ modulate the inflammatory response by secreting pro- and anti-inflammatory factors, such as cytokines, chemokines, matrix metalloproteinases, and growth factors. Furthermore, they phagocytise apoptotic cells, and promote neovascularisation and angiogenesis (Pinto *et al*., 2014; Ma *et al*., 2018; Dick *et al*., 2019). FB, on the other hand, are vital in the remodelling process by secreting various regulatory mediators and by driving the deposition of extracellular matrix (ECM) components to prevent ventricular rupture, thereby ensuring the maintenance of structural and mechanical integrity of the injured heart (Weber *et al*., 2013; Rog-Zielinska *et al*., 2016).

Classically, cardiac cell-type composition has been determined by fluorescence-activated cell sorting (FACS) of immunolabelled cell populations (Banerjee *et al*., 2007). While FACS analysis enables a quantitative analysis of the main cell types of the heart, the approach is biased towards cell-types that survive the applied cell isolation method. Pinto *et al*. partially overcame this limitation by combining FACS counting with immunohistochemical analysis of thin tissue sections, the latter allowing cell type quantification in consecutively cut sections, but with limited 3D information. Using the combined approach, they estimated the percentage of resident cardiac FB and MΦ to account for ∼ 15% and 7% of the NM population, respectively, with the abundance of individual NM populations differing across the various heart chambers (Pinto *et al*., 2016). While state-of-the-art multi-omics approaches allow one to take into account many marker genes for identification of cell subpopulations and their activation states, the sample preparation is associated with similar limitations as apparent for classical approaches. For example, single-cell RNA sequencing relies on prior enzymatic tissue digestion and single-cell sorting, and recently developed fluorescence *in-situ* hybridisation methods such as seqFISH and Merfish are based on fluorescent imaging of thin myocardial tissue sections. Nevertheless, these novel technologies have strongly advanced our understanding of myocardial tissue composition, and dynamics and diversity of NM in specific, both in healthy and remodelled myocardium (Mohenska *et al*., 2022; Anto Michel *et al*., 2022; Kuppe *et al*., 2022; Lother & Kohl, 2023).

In order to better understand cell-type specific functions and heterocellular crosstalk, we need to assess each cell type’s 3D distribution and morphology across the different heart chambers. 3D histology in combination with magnetic resonance imaging represents a powerful approach for quantitative differentiation of volumes occupied by CM and NM (Burton *et al*., 2014), but lacks information on NM subpopulations and the geometry of individual cell networks at the microscale. Genetic labelling by fluorescent reporter proteins (FP) facilitates the distinction and visualisation of tissue-resident FB and MΦ, as well as their subtypes (Acharya *et al*., 2011; Yona *et al*., 2013; Kanisicak *et al*., 2016). However, imaging FP at high resolution in large myocardial volumes has been challenging, mainly due to visible light scattering. Recently developed tissue clearing techniques can be applied to obtain optically transparent heart samples, and labelled cell populations can be transmurally imaged at least in murine hearts (Johnston *et al*., 2017). Tissue clearing is based on removing lipids to reduce light scattering and on matching the refractive index between the tissue and its surrounding solution. Several clearing methods have been applied to the heart; including solvent-based (Three-Dimensional Imaging of Solvent-Cleared Organs, 3DISCO, and Bleaching-Augmented solvent-bAsed Non-toxic Clearing, BALANCE), alcohol dehydration (Clear, Unobstructed Brain Imaging Cocktails, CUBIC), and hydrogel embedding (Clear-Lipid exchanged Acrylamide-hybridized Rigid Imaging/Immunostaining/ In situ hydridization-compatible Tissue-hYdrogel, CLARITY) (Chung *et al*., 2013; Nehrhoff *et al*., 2016; Merz *et al*., 2019; Goodyer *et al*., 2019). The choice of clearing approach depends on the ability to preserve endogeneous fluorescence and architecture of the sample after the clearing process, together with clearing time and compatibility to use immunolabelling to target specific proteins or enhance signals emitted by FP (Tian *et al*., 2021; Olianti *et al*., 2022).

Cardiac application of tissue-clearing techniques includes studies on the distribution of vasculature, immune cells, FB, and neurons, with a focus on lesioned mouse hearts that were most commonly imaged by light sheet microscopy. These studies explored the extent of tissue damage and the 3D cell distribution in cleared hearts post-injury, but did not assess cell morphology, volume, or area. For example, Merz *et al*. studied immune cell infiltration and re-vascularisation at defined time points following ischaemia-reperfusion (I-R) injury using BALANCE-cleared mouse hearts (Merz *et al*., 2019). A drawback of BALANCE is that it may quench endogenous fluorescence. By comparison, the X-CLARITY method was shown to preserve endogenous fluorescence emitted by genetically encoded FP targeted to specific cell types by cre-loxP recombination (Johnston *et al*., 2017). Using this approach, Fischesser *et al*. visualised the dynamic 3D patterns of FB in fibrotic and ischaemic injury models (Fischesser *et al*., 2021). 3D structural information adds value especially when combined with functional experiments and/or computational models. For example, Zhu *et al*. generated maps of sympathetic nerve structure across the whole mouse heart following myocardial infarction, which were aligned with the corresponding maps of electrical propagation obtained by optical voltage mapping using semi-automated algorithms. This allowed them to directly correlate structural changes in sympathetic innervation with altered electrical propagation in infarcted hearts (Zhu *et al*., 2022).

In the here presented work, we applied the X-CLARITY technique to murine hearts selectively expressing Channelrhodopsin-2 (ChR2) fused to the enhanced yellow fluorescent protein (eYFP) in either FB or MΦ. We imaged membrane-bound eYFP fluorescence by confocal microscopy and developed a segmentation algorithm to reconstruct 3D models of FB and MΦ populations at sub-micrometre resolution. The resulting 3D reconstructions were used to determine cell morphology, distribution, surface area, volume and fractional volume of FB and MΦ in all chambers of the healthy mouse heart, as well as in scarred ventricles four-weeks after cryo-ablation. We observed that FB and MΦ have similar morphologies and dimensions across heart chambers, however; FB exist in complex cell networks, whereas MΦ appear primarily as solitary cells. In mature scars following ventricular cryo-ablation, both FB and MΦ numbers are increased, and part of dense and diffusely orientated networks. FB do not only occupy more tissue volume in the scar, but also in remote ventricular regions.

## 2. Methods

All mouse experiments were carried out according to the guidelines stated in Directive 2010/63/EU of the European Parliament on the protection of animals used for scientific purposes and were approved by the local authorities in Baden-Württemberg (G16-131,G20-001, and G22-090).

### Animal model

Heterozygous mice (C57BL/6J) transgenic for Cre recombinase fused to two modified estrogen receptor ligand binding domains under the control of the transcription factor 21 promoter (Tcf21-MerCreMer (Acharya *et al*., 2011)), or transgenic for Cre recombinase fused to a single mutant estrogen ligand binding domain under the control of the chemokine receptor 1 promoter (Cx3cr1-CreERT (Yona *et al*., 2013)) were crossed with homozygous mice transgenic for floxed Cop4 H134R-eYFP (B6; 129S-Gt(ROSA)26Sortm32(CAG-COP4*H134R/EYFP)Hze/J (Madisen *et al*., 2012)). The resulting offspring expressed ChR2-eYFP in either FB (Tcf21-ChR2) or MΦ (Cx3cr1-ChR2) upon tamoxifen induction (Fernández *et al*., 2021). Mice were weaned at P28–35 and ear snips were taken for PCR-based genotyping. For assessment of healthy hearts, double-positive mice received daily intraperitoneal injections of Tamoxifen at 0.11 mg per g body weight for 5 days at 8 weeks of age. Experiments on hearts from healthy mice were performed 14-21 days later (i.e. at an age of 10-12 weeks)

### Ventricular cryoablation

Left ventricular cryoablation was applied on 10-week old double-positive transgenic mice, as previously described (Simon-Chica *et al*., 2023*a*). Tcf21-ChR2 mice received Tamoxifen injections starting 14 days before the day of surgery as outlined above, while Cx3cr1-ChR2 mice received tamoxifen starting on day 7 after surgery (see supplementary figure 1). Before the surgical procedure, mice received a subcutaneous injection of 250 µL analgesia solution (10 µg/mL buprenorphine [Temgesic, Indivior Inc., North Chesterfield, VA, USA] in 154 mM NaCl [0.9% m/V, B. Braun Melsungen, Melsungen, Germany]) into the scruff, followed by an intraperitoneal (i.p.) injection of 80-100 µL anaesthesia solution (20 mg/mL Ketamine [Ketaset, Zoetis, Parsippany-Troy Hills, NJ, USA], 1.4 mg/mL Xylacin hydrochloride [0.12% Rompun, Bayer, Leverkusen, Germany], in 114 mM NaCl [0.67% m/V, Rompun]). For volume substitution of the expected perioperative blood and evaporative fluid loss, 500 µL glucose solution (278 mM glucose = 5% (m/V), B. Braun Melsungen) was injected i.p. An eye ointment (Bepanthen containing 50 mg/mL dexpanthenol, Bayer) was applied, and the chest and neck of animals were disinfected using Softasept N (B. Braun Melsungen). The left side of the thorax (precordial region) and the right leg were shaved. Mice were placed on a warming platform of a small animal physiology monitoring system (Harvard Apparatus, Holliston, MA, USA), and the front extremities were fixed with tape. During the surgical procedure, the body temperature of the mouse was monitored using a rectal thermometer and maintained between 36 and 37 °C. Thermometer and tail were fixed with tape. After, the orotracheal intubation, the mouse was connected to an external ventilation system (Kent Scientific, Torrington, CT, USA; 40% O_2_, 120 breathing cycles per minute). Isoflurane was applied at 5% until the animal stopped spontaneous respiratory movements, and then reduced to 2.0-2.5%. An infrared blood oximeter was attached to the right leg to follow hemoglobin oxygen saturation. A lateral thoracotomy was achieved by cutting the skin and muscles along the third intercostal space. After introducing a rib spreader, the pericardium was cut and the epicardial surface of the free wall of the left ventricle was carefully dried-blotted using a cellulose pad. A metal probe (stainless steel, 2.5-mm hexagon) pre-chilled in liquid nitrogen was applied for 8-10 s on the free left-ventricular mid-wall, avoiding major coronary vessels. After retraction of the probe, the time until the tissue regained deep-red colour was monitored (typically within 5 to 10 s). The rib spreader was removed and the thorax closed using a 6-0 silk suture around the third and fourth rib (4-5 single knots). Before final closure, remaining air was removed from the thorax with a small cannula. Isoflurane application was stopped and the skin was closed with a 6-0 silk suture. Once the mouse started breathing independently, intubation and fixation were terminated, and the mouse was transferred to a heated and oxygenated wake-up chamber. Analgesia was maintained for 72 h post-ablation by twice-daily subcutaneous injection of 250 µL of 10 µg/mL buprenorphine (in 154 mM NaCl; morning and late afternoon). During the night, buprenorphine was supplied via the drinking water (10 µg/mL buprenorphine [Subutex lingual tablets, Indivior] in 20 mM glucose solution). Hearts were harvested for X-CLARITY experiments 28 days after surgery.

### Tissue Processing

Mice were euthanised by cervical dislocation, their chest opened, and heart removed. Hearts were immediately washed in warm (37 °C) heparin-containing (5 unit/mL) physiological saline solution (containing in mM: 140 NaCl, 6 KCl, 10 HEPES, 10 glucose, 1.8 CaCl_2_, 1 MgCl_2_, pH 7.4 [adjusted with NaOH], 300±5 mOsmol L^-1^) to remove blood before placing the heart in cold (4 °C) solution to arrest beating. The aorta was cannulated and the heart was Langendorff-perfused with 10 mL of physiological saline solution (as above, but without heparin), followed by 10 mL 4% (m/V) paraformaldehyde in phosphate buffered saline (PBS [Sigma Aldrich, St Louis, MO, USA], containing in mM: KH_2_PO_4_ 136, NaCl 58, Na_2_HPO_4_-7H_2_O 268). Hearts were kept in this solution for 5 h at room temperature, then washed three times with PBS, and stored in PBS overnight at 4 °C. The PFA-preserved heart was dissected into the following parts: (1) the atria in one piece, (2) right ventricle (RV) together with the septum, and (3) the left ventricular free wall (LV) before starting the clearing procedure (X-CLARITY, Logos Biosystems, Anyang, South Korea). The applied clearing protocol was based on the manufacturer’s instructions for mouse brains, but with adjusted parameters for the electrophoresis step. Tissue pieces were incubated with the hydrogel solution (Logos Biosystem) containing 4% acrylamide and a polymerisation initiator containing 0.25% Azobis [2-(2-imidazolin-2yl)propane] dihydrochloride used to create the polyacrylamide gel matrix (Logos Biosystems) at 4°C overnight, and polymerised using the X-CLARITY polymerisation system for 3 h at 37 °C while applying −90 kPa via a vacuum pump. Afterwards the tissue was agitated for 1 min, and rinsed with electrophoretic tissue clearing (ETC) solution containing 200 mM sodium borate buffer and 4% sodium dodecyl sulphate (SDS) at pH 8.5 (Logos Biosystems), before being incubated with ETC solution in the X-CLARITY tissue clearing system for 2 h at 37 °C, while applying a current of 0.8 A. Finally, hearts were washed and incubated in 1% Triton X-100 (Sigma Aldrich) in PBS for 24 h at 37 °C to completely remove SDS.

### Immunohistochemical staining of cleared tissue

Cleared tissue from Tcf21-ChR2 hearts were incubated under continuous agitation with Hoechst 33342 (Ab228551 at 1:500 in PBS, Abcam, Cambridge, UK) overnight at 37 °C, and washed three times with PBS for 20 min at room temperature. Tissues from Cx3cr1-ChR2 hearts were additionally immunostained to amplify the eYFP fluorescence. Tissues were incubated with blocking solution (0.5% Triton X, 2.5% bovine serum albumin [Sigma-Aldrich], 2.5% donkey serum [Sigma-Aldrich]) for one day at 37 °C before being incubated with the primary antibody at 1:500 for 3 days at 37 °C. Then, tissues were washed three times by incubation with PBS containing 0.1*%* Tween for 4 h at 37 °C under continuous agitation, followed by incubation with the secondary antibody, donkey anti-goat Alexa Fluor 488 (Z-25006, Life Technologies, Carlsbad, CA, USA) at 1:1000 for 3 days at 37 °C. Unbound secondary antibody was removed by three washing steps in PBS containing 0.1 % Tween for 4 hours each. The nuclei staining with Hoechst 33342 was performed as described for Tcf21-ChR2 hearts. Stained tissues can be stored in PBS at room temperature for a maximum of two days.

### Confocal microscopy

Prior to confocal imaging, cleared and stained heart tissues were briefly washed with distilled water and incubated with X-CLARITY mounting solution (refractive index of 1.46; Logos Biosystems) for 5 h at room temperature. Hearts were transferred to an 8-well μ-dish with glass-bottom (ibidi, Gräfelfing, Germany) containing X-CLARITY mounting solution. Image stacks was recorded with an inverted confocal microscope (TCS SP8 X, Leica Microsystems, Wetzlar, Germany) either with a water immersion objective (HC PL APO 40x/1.10) or a glycerol immersion objective (HC PL APO 63x/1.30). Alexa Fluor 488 and eYFP were excited by the 488-nm and 514-nm “lines” of a white-light laser, respectively. Hoechst 33342 was excited using a 405 nm diode laser. Fluorescence signals were recorded with sensitive hybrid detectors or a photomultiplier detector for Hoechst 33342 and suitable spectral detection windows. Imaging was performed using the Leica Application Suite X (LAS-X) with the LIGHTNING module, which enables deconvolution-based super-resolution imaging by optimizing the imaging settings for deconvolution and computing the deconvolved 3D image.

### Quantification of ChR2-eYFP expression in FB and MΦ in 2D

The fractional area occupied by FB and MΦ in healthy tissue was calculated using the membrane-bound eYFP signals of 2D overview images, providing an estimate on the distribution and relative density of NM in tissue. To do this, we created an automated segmentation algorithm, which generates masks of the areas occupied by eYFP-positive cells and by cardiac tissue. Raw images were loaded into numpy arrays using the tifffile library. All images were de-noised by applying a Gaussian filter with σ =1 pixel. The eYFP-positive area (foreground) and the tissue area (background) were determined from the intensity histogram of filtered images. All pixels with an intensity below D + 0.5·SD (D denotes the mode and SD the standard deviation of the intensities) were used to define the background, and all pixels above D + 3.5·SD were defined as foreground for MΦ, and all pixels above D+ 1·SD were labelled as foreground for FB. Following binary segmentation, the tissue mask was further processed by two iterations of binary opening followed by five iterations of binary closing. The eYFP masks of MΦ and FB were further processed with five iterations of binary dilation, followed by three iterations of binary erosion. Objects with fewer than 300 pixels were considered noise and were removed. The fractional area of labelled FB and MΦ was calculated from the ratio between foreground and background masks. MΦ numbers were counted using a morphology module to label the mask objects.

### 3D reconstructions of FB and MΦ

We assessed the 3D morphology, distribution, surface area, volume, and fractional volume of FB and MΦ in healthy and cryo-ablated hearts. To do this, we recorded z-stacks of eYFP fluorescence (FB) and Alexa-Fluor 488 (MΦ) by confocal imaging of cleared heart chambers. For healthy hearts we imaged the left and right atria (LA and RA), the LV, RV, and the septum, whereas for cryo-ablated hearts, we imaged the scar centre, the border zone and the remote myocardium of the LV and RV. 3D reconstructions of FB and MΦ were obtained with a semi-automatic algorithm written in Python, previously used by Simon-Chica et al. for determination of MΦ dimensions in LV tissue (Simon-Chica *et al*., 2022).

Raw images were loaded into numpy arrays using the tifffile library. Each channel was filtered with a Gaussian filter kernel (σ = 1 voxel) using the ndimage module from SciPy. Background was removed from each channel by subtracting the mode of the intensity histogram from each voxel, and by setting any resulting negative values to 0. Depth-dependent attenuation was corrected in each channel by fitting an exponential function to the average intensities of each slice, and by scaling the intensities by the inverse of the resulting function. Next, binary segmentations were created using a threshold (T), based on the average (A) and SD of the intensity per channel: T = A + f_SD_ · SD for FB or MΦ-specific fluorescence, and T = A + 2 SD for nuclei stain. f_SD_ was manually chosen between 1.75 and 2, based on visual inspection of image quality. Five iterations of binary dilation, followed by five iterations of binary closing were performed on FB or MΦ segmentations using an ellipsoidal structural element with axis length of 2 voxels in x- and y-directions and 1 voxel in z-direction. Four iterations of binary erosion were performed on each slice for nuclei segmentation, followed by three iterations of binary closing, using the same structuring element as for FB and MΦ.

Contiguous objects in the binary FB, MΦ, and nuclei maps were uniquely labelled, and objects containing less than 2,000 voxels were removed to eliminate small staining artifacts. Voxel sizes of 3D reconstruction from *in situ* FB and MΦ are shown in supplementary tables 1 and 2. The scikit-image library was used for these operations (Van Der Walt *et al*., 2014). Objects in FB and MΦ segmentations touching stack boundaries were removed to exclude incompletely imaged cells in later analyses.

For each of the segmented nuclei, co-localisation with any cell mask was determined. If more than Z % of voxels of a segmented nucleus were co-localised with a segmented cell, the nucleus was considered belonging to that cell. Z was chosen manually between 5 and 15, based on visual assessment of the nuclei segmentations. The low percentage results from the fact that voxel co-localisation only occurs at the surface of nuclei segmentations – due to eYFP expression in the cell membrane rather than the nucleus, thus FB or MΦ masks have “holes” where nuclei are located. Cell masks co-localised with nuclei segmentations were merged and binarised, as nuclei could theoretically connect previously unconnected objects.

A Gaussian filter with σ = 2 voxels in x- and y-directions and 0.5 voxels in z-direction was applied on the binarised segmentation, followed by one iteration of grayscale dilation with a box structuring element of 15×15×5 voxels. This created monotonically decreasing intensities from the inside towards the outside of each cell that could be used to adjust over-/ under-segmentation when creating 3D surfaces. An iso-surface threshold, manually chosen based on image quality between 80 and 180 (out of 255), was used when computing surfaces, using a marching cubes algorithm from scikit (Lewiner *et al*., 2003). The resulting triangular mesh was further processed and analysed using the open-source Visualisation Toolkit (Schroeder *et al*., 2006). A smoothing filter (vtkSmoothPolyDataFilter) was applied to the mesh with 3,000 iterations, a convergence threshold of 0.1, and a feature angle of 90°. Volumes and surface areas of each of the resulting 3D reconstructions were computed using the vtkMassProperties algorithm.

### Statistical analysis

Data was compared between groups using a linear mixed effects model with fixed effects for cell type and chamber, and random effects for mouse and sample. An ANOVA test was applied to the fixed-effect coefficients and p-values <0.05 were considered as statistically significant. We compared the differences in surface area, volume, and the percentage of the fractional volume between MΦ and FB in the different chambers of the healthy hearts. Additionally, we used this model to compare the fractional volume of FB and MΦ between remote and scar areas in injured hearts and healthy hearts from LV. For analysis of healthy hearts, a total of 54 regions of interest (ROIs) was analysed including the left atrium (LA), right atrium (RA), LV, RV, and septum from Tcf21-ChR2 hearts (N= 3 hearts). In addition, 30 ROIs were included from the different chambers of Cx3cr1-ChR2 hearts (N= 3 hearts). For cryo-ablated Tcf21-ChR2 hearts, 17 ROIs from remote areas from LV and RV were compared to 14 scars areas (N=3 hearts). For cryo-ablated Cx3cr1-ChR2 hearts, we analysed 10 ROIs both from remote myocardium and from the scar (N=3 hearts).

## 3. Results

### Distribution of resident FB and MΦ in 2D in healthy hearts

To assess the overall abundance and distribution of FB and MΦ in healthy hearts, we imaged cleared heart chambers with membrane-bound eYFP expression in either FB or MΦ, respectively. Heart chambers were optically cleared using an adapted X-CLARITY approach and incubated in refractive index matching solution before confocal microscopy. While this did not yield completely optically transparent samples, we could detect eYFP fluorescence at imaging depths up to 500-800 μm. We took overview images of a cross-section of each of the LV, LA and RA (Figures 1 and 2), and we quantified the percentage of the eYFP-positive area per tissue area using an automated analysis algorithm.

**Figure 1.**
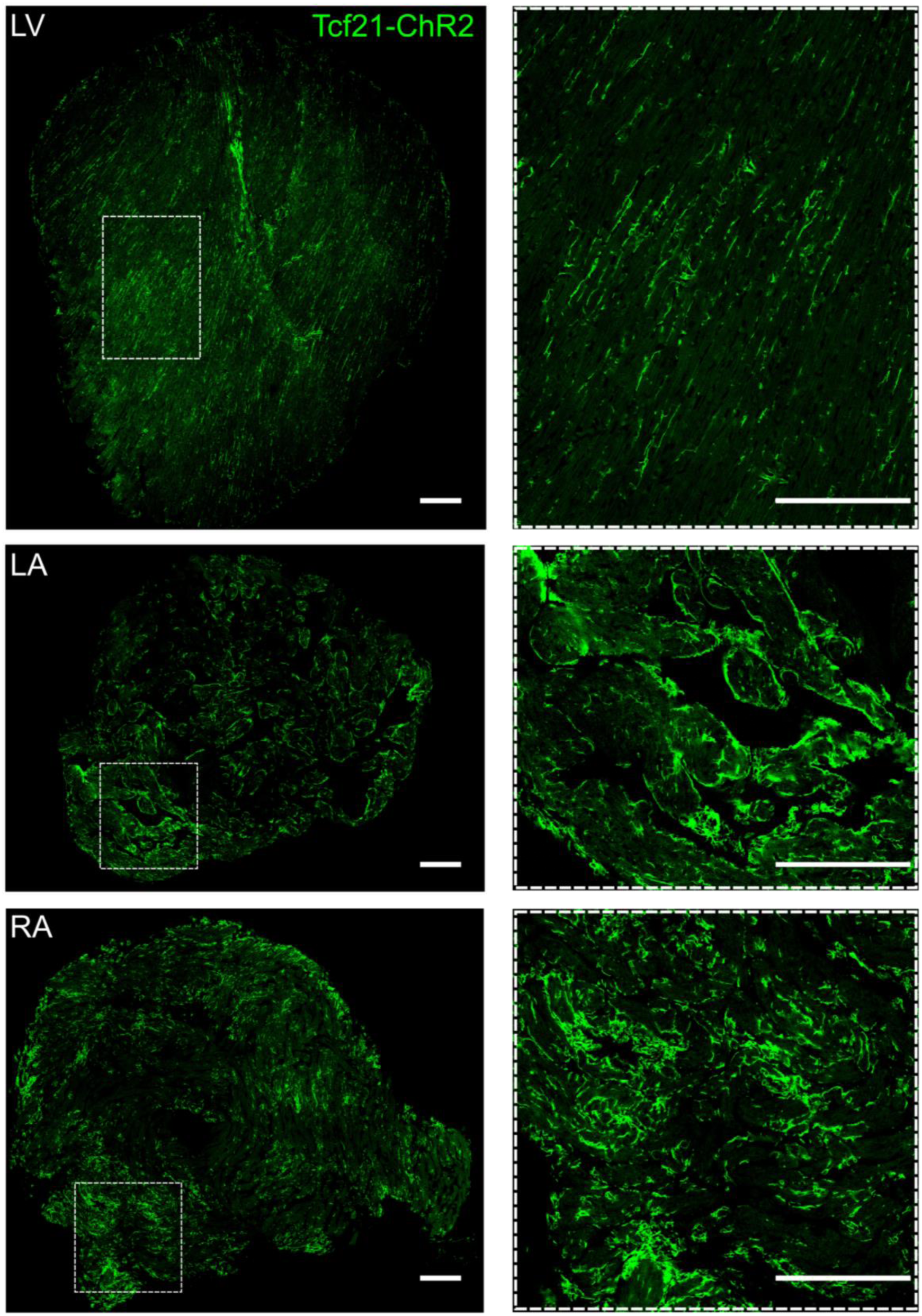
2D overviews of FB distribution in LV, LA, and RA of cleared Tcf21-ChR2 heart. Confocal images of FB expressing ChR2-eYFP in different chambers of the heart (scale bars – 500 μm). Green – eYFP fluorescence.

**Figure 2.**
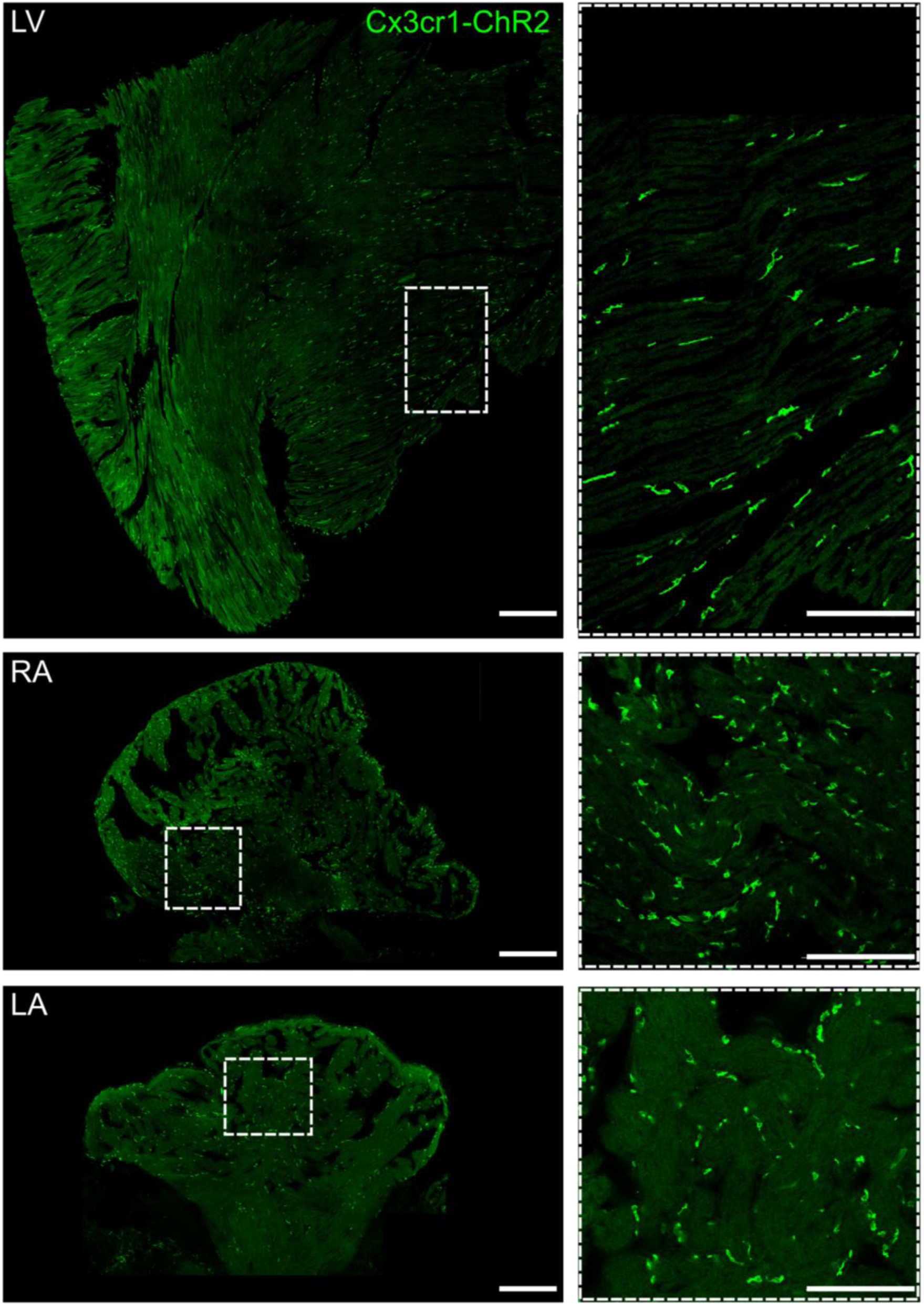
2D overviews of MΦ distribution in LV, LA and, RA of cleared Cx3cr1-ChR2 heart. Confocal images of MΦ labelled with anti-GFP antibody to amplify the eYFP signal (scale bars – 500 μm). Green – Alexa488 fluorescence.

Figure 1 shows representative overview images from a heart expressing eYFP in FB. FB are abundant in the LV and in the atria, where they form extended, interconnected cell networks. In the LV, FB networks have longitudinal patterns following the regular alignment of CM (long axis of CM parallel to the LV surface and thus to the image plane), as recognisable from the tissue autofluorescence. In contrast, FB observed in LA and RA form networks that show both longitudinal and torturous patterns, in line with a less regular alignment of atrial CM and with the orientation of the image plane sectioning CM at different angles (both parallel and perpendicular to their long axis). For the shown representative overview images, we calculated the percentages of the fractional area of eYFP-expressing FB in respect to total tissue area to be 4.2% in the LV, 8.8% in the LA and 10.3% in the RA (see supplementary table 3 for details).

Figure 2 shows representative overview images from a heart expressing eYFP specifically in MΦ. Similar to FB, MΦ were abundantly found in the LV, the LA and the RA, but in contrast to FB, MΦ appeared mainly as solitary cells. Based on their solitary appearance, absolute MΦ numbers can be quantified in images from the different heart chambers. In the analyzed 2D images, we found a higher number of MΦ per tissue area in the RA in comparison to the LA and the LV. In the representative images, the fractional area of MΦ per tissue area (supplementary table 3) were 0.7 % in LV, 0.8 % in the LA, and 1.5% in the RA. In representative 2D images, we observed FB to be more interconnected and occupying larger fractional areas compared to tissue-resident MΦ in the LV, RA, and LA of healthy murine hearts.

### 3D reconstructions of resident FB and MΦ in healthy mouse hearts

In order to quantitatively assess the abundance, dimensions and interconnectivity of NM across the different heart chambers, we reconstructed the 3D structure of eYFP-labelled FB and MΦ in cleared heart tissue. While Hoechst stained nuclei independent of cell type (both CM and NM, see image projections on the left in Figure 3 and 4), we developed an algorithm for identifying nuclei within labelled NM (networks), based on co-localisation between eYFP and Hoechst fluorescence signals (see methods for details).

**Figure 3.**
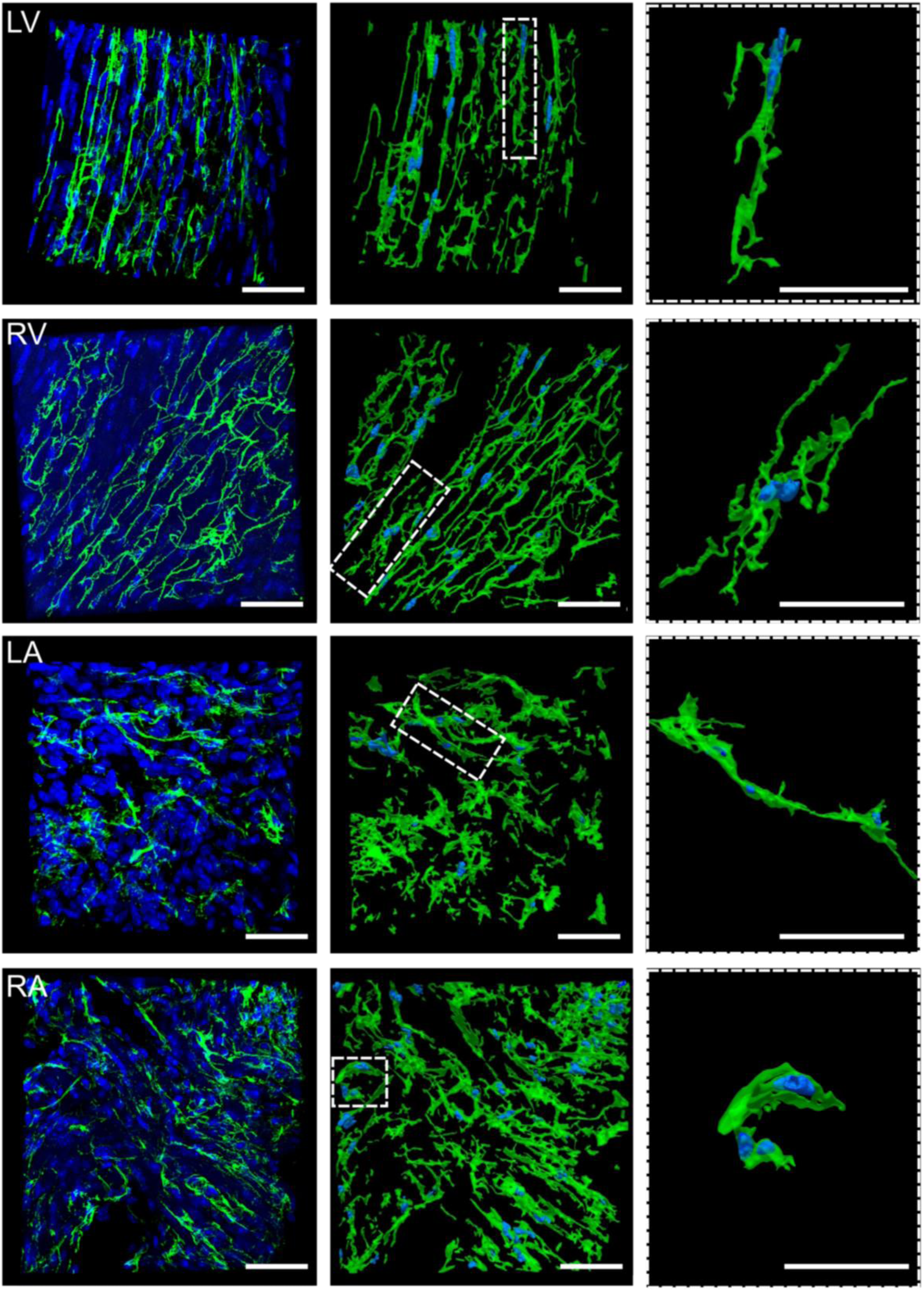
3D projections and corresponding reconstructions of ChR2-eYFP-labelled FB in Tcf21-ChR2 cleared heart. 3D projection of confocal image stack (left) and 3D reconstructions (middle, right) of ChR2-eYFP expressing FB in LV, RV, LA and RA (scale bars – 50 μm). Green – eYFP; blue – nuclei staining (Hoechst).

**Figure 4.**
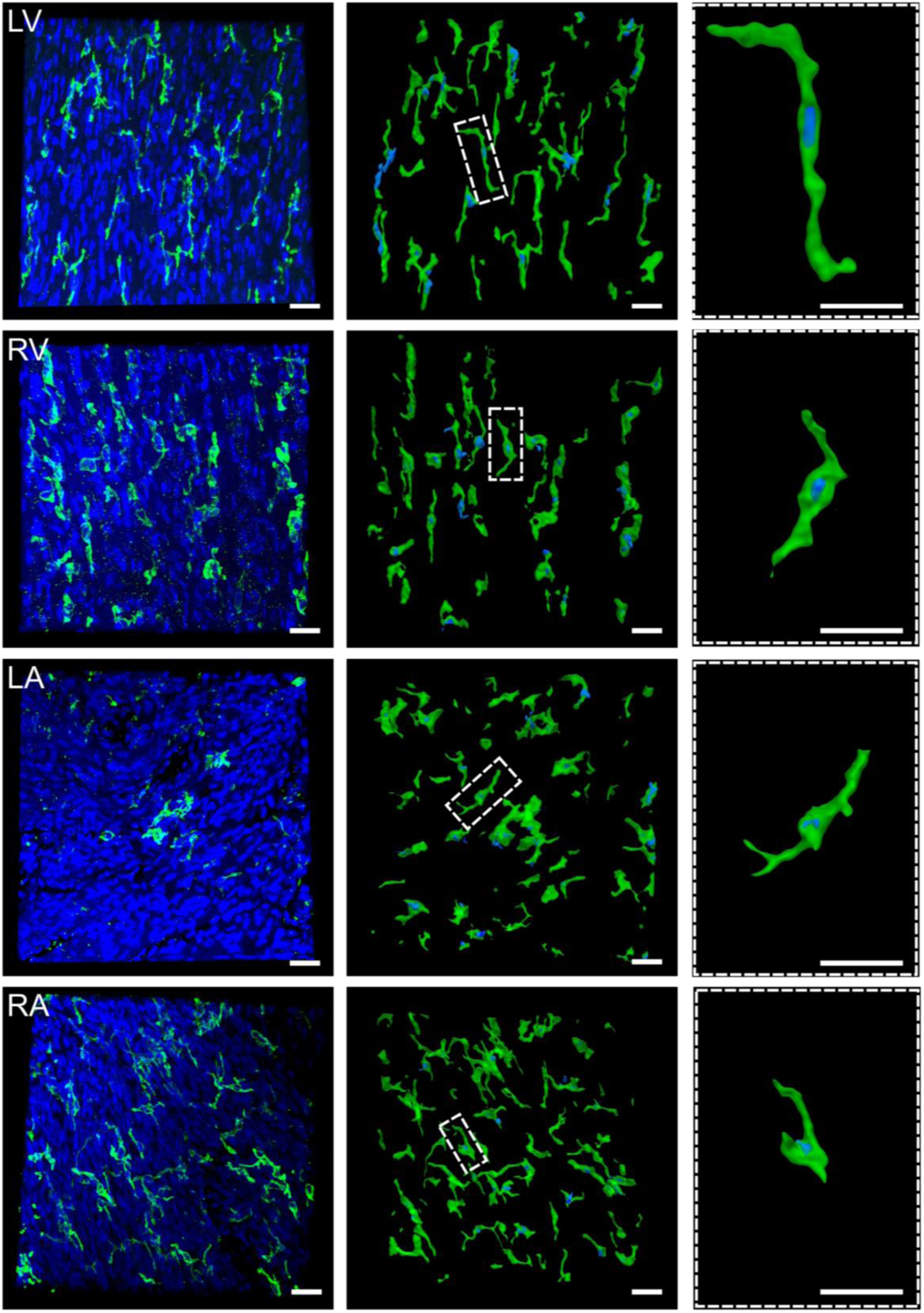
3D projections and corresponding reconstructions of ChR2-eYFP-labelled FB in Cx3cr1-ChR2 cleared heart. 3D projection of confocal image stack (left) and 3D reconstructions (middle, right) of ChR2-eYFP expressing MΦ in LV, RV, LA and RA (scale bars – 50 μm). Green – Alexa488 fluorescence; blue – nuclei staining (Hoechst).

Figure 3 shows representative tissue volumes containing eYFP-labelled FB and stained nuclei in LV, RV, LA, and RA. Panels on the left show 3D projections of confocal image stacks, next to the corresponding 3D reconstructions (middle) and higher magnification images depicting selected individual cells (right panels). Consecutively imaged planes from z-stacks and the corresponding 3D reconstructions of FB in LV are visualised also in supplementary video 1.

In line with observations in 2D, 3D reconstructions reveal different distributions and morphologies of FB in individual heart chambers, especially when comparing ventricular to atrial FB. In LV and RV (Figure 3A and B), FB follow the regular 3D alignment of CM and are characterised by longitudinal (parallel to predominant CM orientation) and transversal (perpendicular to CM orientation) extensions, thereby forming extended networks within the imaged tissue volumes. In fact, defining the borders of individual FB is nearly impossible, even in healthy ventricular tissue. In the atria (Figure 3C and D), FB also form dense, interconnected networks, but in contrast to the ventricles, network structure is less regular and FB adopt both tortuous and sheet-like morphologies not commonly observed in the ventricles.

Figure 4 shows 3D projections of confocal image stacks (left panel) of eYFP-expressing MΦ and labelled nuclei next to 3D reconstructions of nuclei-containing MΦ (middle and right). Similar to FB, and in contrast to circulating leukocytes, tissue-resident MΦ are elongated cells characterised by tree-like extensions from the nuclei and smaller protrusions, following the structure of CM. MΦ orientation (long-axis) follows longitudinal CM alignment in the ventricles, while MΦ are less regularly aligned in the atria. In contrast to FB, MΦ exist predominantly as solitary, *i.e.* not interconnected cells, across all chambers of the healthy mouse heart (for LV, see supplementary video 2). While most MΦ were solitary, occasional small MΦ clusters surrounding CM were found.

### Quantitative analysis of FB and MΦ dimensions and abundance in 3D across all heart chambers

Based on the solitary nature of most MΦ, reconstructions of isolated, mononuclear MΦ were used for quantifying MΦ surface area and volume in healthy myocardium (for measuring the fractional volume, we used all segmented MΦ including small cell clusters containing multiple nuclei). The surface area and volume per FB were determined by dividing the dimensions of FB networks by the number of co-localised nuclei. Figure 5A and B compare the average surface area and volume of FB in networks (dimensions per nuclei) to the surface area and volume of individual, mononucleated MΦ. The numerical values for the individual heart chambers are listed in table 1. Most importantly, we found that FB and MΦ had similar dimensions across all chambers of the heart. There was no difference in cell volume between the two cell types, across all chambers (p=0.648). However, there were significant differences in surface area between FB and MΦ in the atria: FB showed larger average surface areas in the LA (p=0.0168) and in the RA (p=0.0041), when compared to MΦ in the same chambers. Looking into the individual cell populations, the surface area of FB was significantly smaller in LV tissue compared to LA (p=0.0292) and RA (p=0.0251). Sizes of reconstructed MΦ were not different between the atria and ventricles.

**Figure 5.**
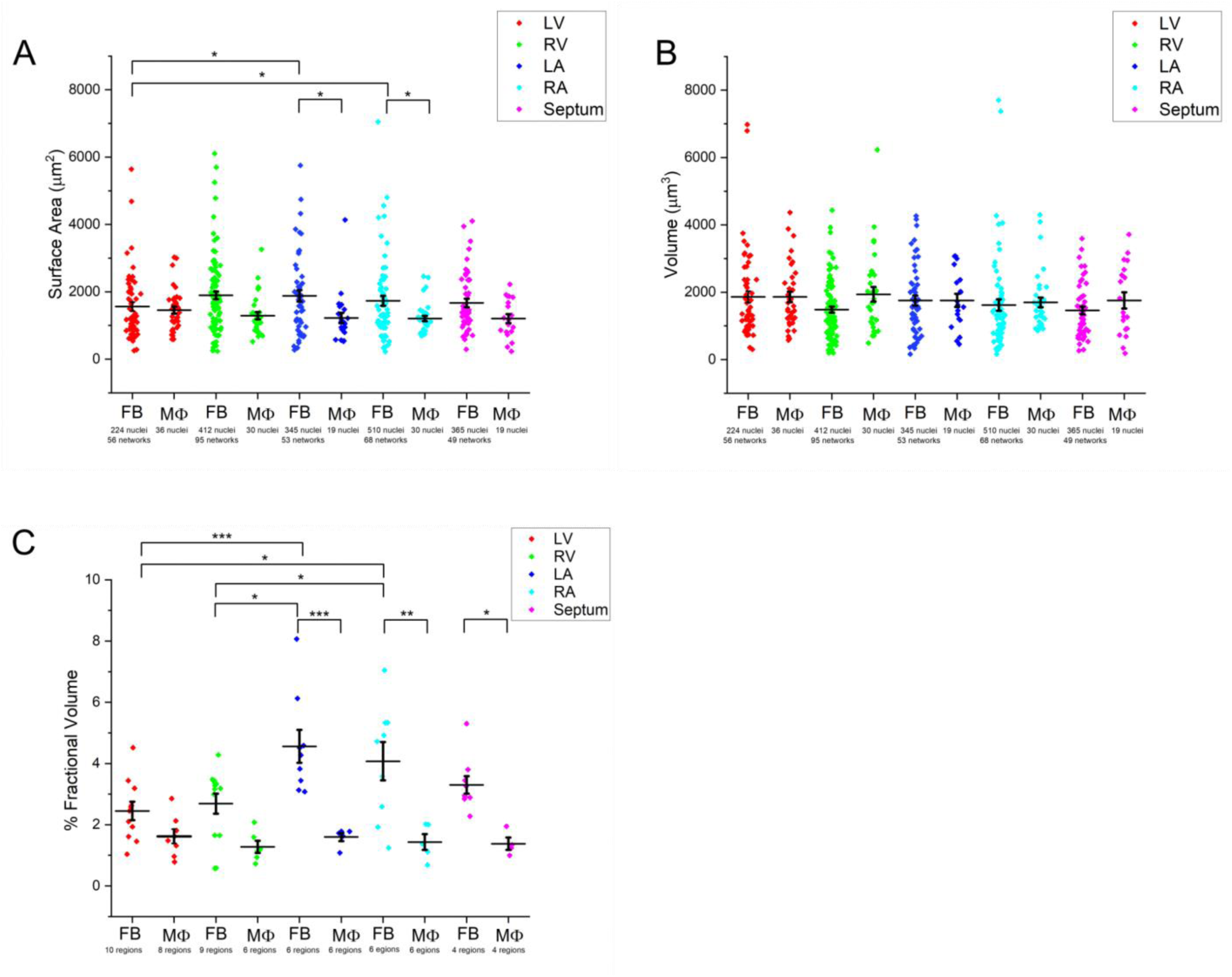
Quantitative comparison of surface area, volume and fractional volume of FB and MΦ populations in healthy cleared hearts. Surface area (A) and volume (B) of FB and MΦ across the cardiac chambers. FB from LV (n=56 networks), RV (n=95 networks), LA (n=51 networks), RA (n=68 networks), and septum (n=49 networks) in N=3 Tcf21-ChR2 mouse hearts. MΦ from LV (n=36 cells), RV (n=31 cells), LA (n=19 cells), RA (n=35 cells), and septum (n=19 cells) in N= 3 Cx3cr1-ChR2 mouse hearts. C) Comparison of the percentage fractional volume between FB and MΦ across heart chambers. *P < 0.05, **P < 0.01, ***P< 0.001.

**Table 1.** Average surface area and volume of ChR2-expressing FB and MΦ in healthy mouse hearts. SE – standard error of mean, N – number of FB networks.

| Chambers | FB |  |  |  | MΦ |  |  |
| --- | --- | --- | --- | --- | --- | --- | --- |
| | FB Networks | Nuclei | Surface Area ( $\mu\text{m}^2$) | Volume ( $\mu\text{m}^3$) | Nuclei | Surface Area ( $\mu\text{m}^2$) | Volume ( $\mu\text{m}^3$) |
| **LV** | 56 | 224 | 1560±130 | 1860±170 | 36 | 1460±100 | 1860±160 |
| **RV** | 95 | 412 | 1890±120 | 1480±90 | 31 | 1290±110 | 1940±220 |
| **LA** | 51 | 345 | 1880±210 | 1750±150 | 19 | 1120±100 | 1760±190 |
| **RA** | 68 | 510 | 1840±180 | 1620±170 | 35 | 1210±80 | 1700±150 |
| **Septum** | 49 | 365 | 1670±130 | 1460±120 | 19 | 1200±130 | 1760±240 |

The fractional volume (FV) of FB and MΦ in cleared hearts was calculated by the ratio between the volume occupied by reconstructed cells and the total tissue volume (see table 2). Figure 5C compares the FV occupied by FB and MΦ for each heart chamber. There was a significant difference between the FV of FB and MΦ in the LA (4.6±0.5% *vs* 1.6±0.1%; p= 2.14e-5), RA (4.1±0.6% *vs* 1.4±0.3%; p=0.0014) and septum (3.3±0.3% *vs* 1.4±0.2%; p=0.0422), where FB occupy larger volume fractions than MΦ. When focusing on FB, we found that FB occupied greater FV in the atria compared to the ventricles. Specifically, volume occupied in the LA (4.6±0.5%; p=4.71e-4 *vs* LV; p= 0.0215 *vs* RV) and in the RA (4.1±0.6%; p=0.012 *vs* LV; p= 0.0154 *vs* RV) was larger than volume occupied by FB in the LV (2.4±0.3%) and the RV (2.7±0.3%), respectively. FV of MΦ was not significantly different between the individual heart chambers (p= 0.622).

**Table 2.**
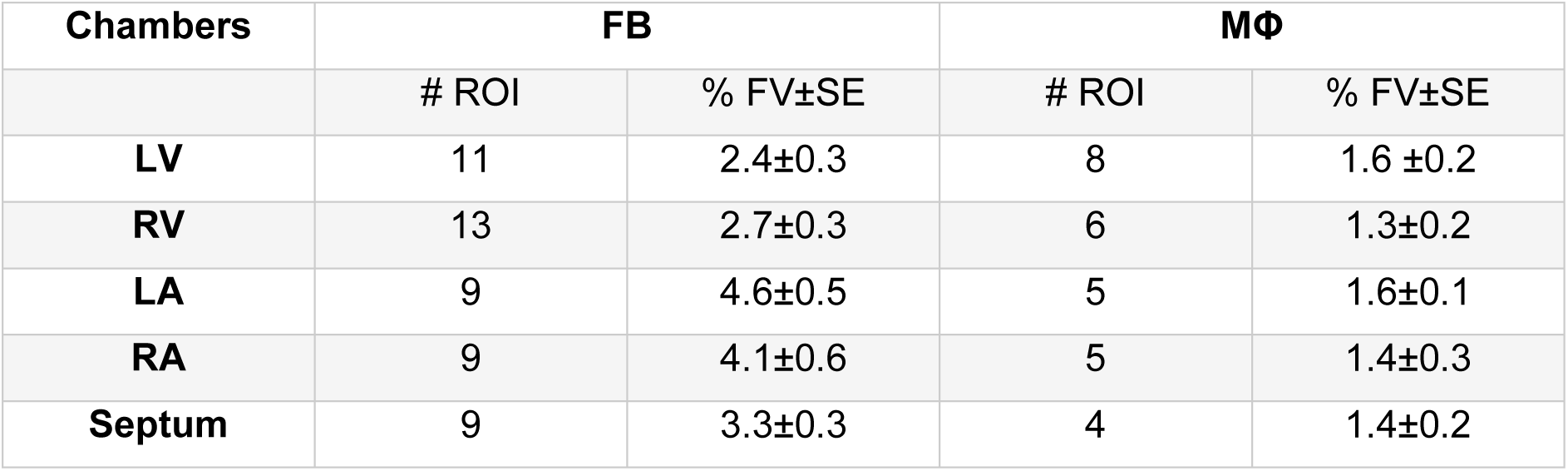
Fractional volume of FB and MФ across heart chambers. SE – standard error of mean.

| Chambers | FB |  | MΦ |  |
| --- | --- | --- | --- | --- |
|  | # ROI | % FV±SE | # ROI | % FV±SE |
| **LV** | 11 | 2.4±0.3 | 8 | 1.6 ±0.2 |
| **RV** | 13 | 2.7±0.3 | 6 | 1.3±0.2 |
| **LA** | 9 | 4.6±0.5 | 5 | 1.6±0.1 |
| **RA** | 9 | 4.1±0.6 | 5 | 1.4±0.3 |
| **Septum** | 9 | 3.3±0.3 | 4 | 1.4±0.2 |

### 3D reconstruction of FB and MΦ in murine hearts after ventricular cryo-injury

We studied the morphology, distribution, and FV of FB and MΦ in murine hearts four weeks after ventricular cryo-injury. Figure 6A and B show the 3D projections of confocal image stacks (left panels) and the corresponding reconstructions of FB networks and nuclei (right panels) in remote myocardium and scar centre of cryo-ablated hearts, respectively (scar ROI, see supplementary video 3). FB networks in remote LV tissue showed a regular, longitudinal alignment in respect to CM orientation, similar to FB networks in healthy LV. In the scar, however, both the total number of nuclei as well as the number of nuclei within FB networks are strongly increased, preventing the reconstruction of individual nuclei within dense cell clusters. In the scar, FB form extended and dense networks, with sheet-like morphologies instead of thin cellular extensions. In accordance, the FV of FB in the scar region was significantly larger than the FV occupied by FB in remote regions (13±1% versus 7.9±0.4%, respectively; p=4.256e-7; Figure 6c). In addition, the FV of FB in remote tissue was larger than their FV in healthy hearts (2.6±0.3%; p=1.644e-6).

**Figure 6.**
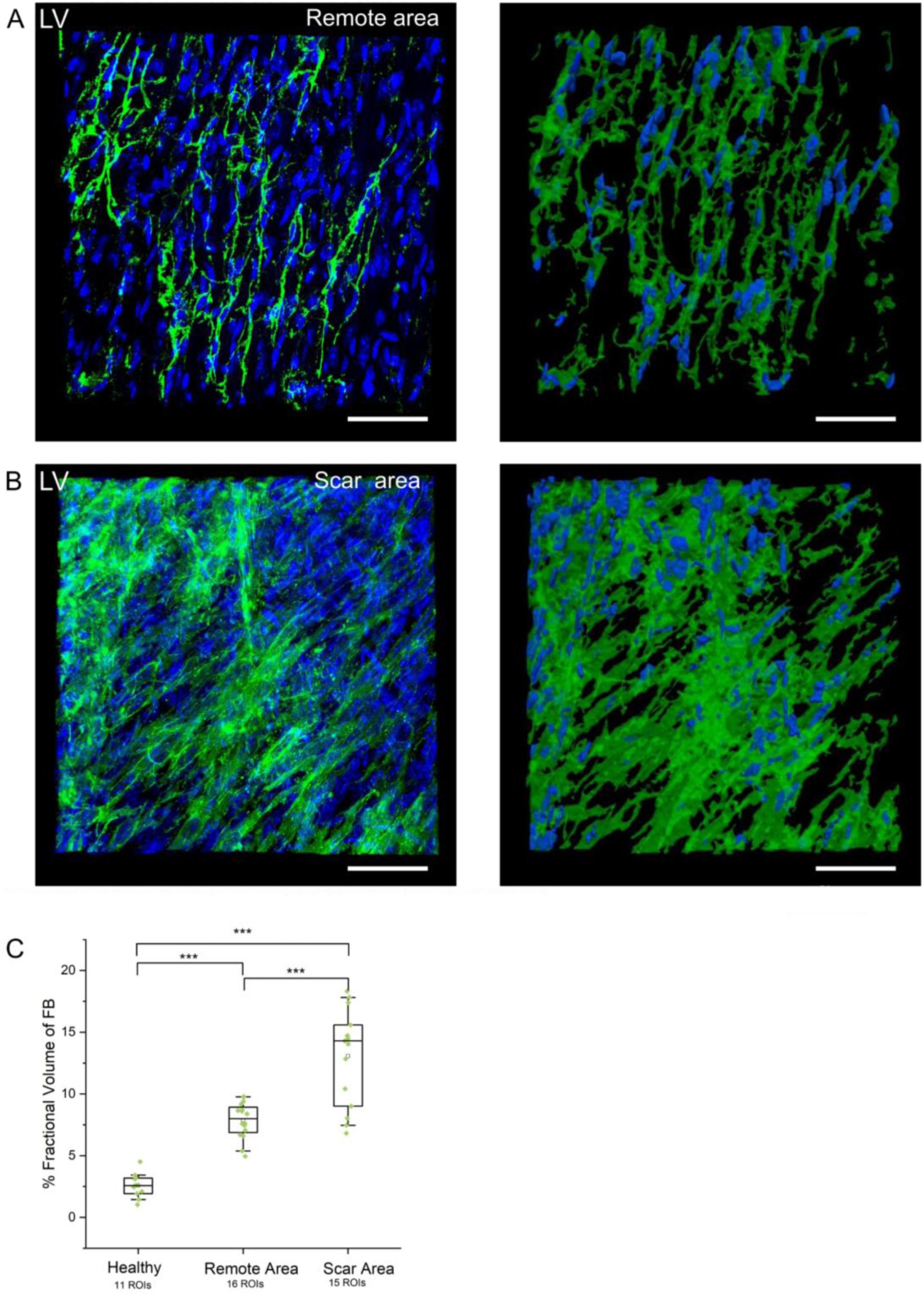
3D projections and reconstructions of FB in LV of cryo-ablated heart. 3D projection of confocal image stack (left) and 3D reconstruction (right) of ChR2-eYFP expressing FB in remote (A) and scar (B) regions of cryo-ablated Tcf21-ChR2 heart (scale bars – 50 μm). C) Fractional volume occupied by FB in LV regions of healthy hearts (n=11 ROI, N= 3 Tcf21-ChR2 hearts), compared to remote (n=16 ROI) and scar (n=15 ROI) regions of cryo-ablated hearts (N=3 Tcf21-ChR2 cryo-ablated hearts).

Figure 7 shows the 3D projections and corresponding reconstructions of MΦ, comparing remote myocardium and scar of a cryo-ablated heart. In remote LV, MΦ show a similar distribution compared to healthy myocardium, and predominantly appear as solitary cells (Figure 7A, see supplementary video 4). Within the scar, however, MΦ are interconnected, forming cell networks that occupy significantly larger FV (7.7±0.6 % in scar compared to 2±0.3% in the remote region; p=2.576e-11). In contrast to observations on FB, FV of MΦ was not different between remote regions of cryo-ablated hearts and healthy hearts (1.6±0.2%; p=0.578).

**Figure 7.**
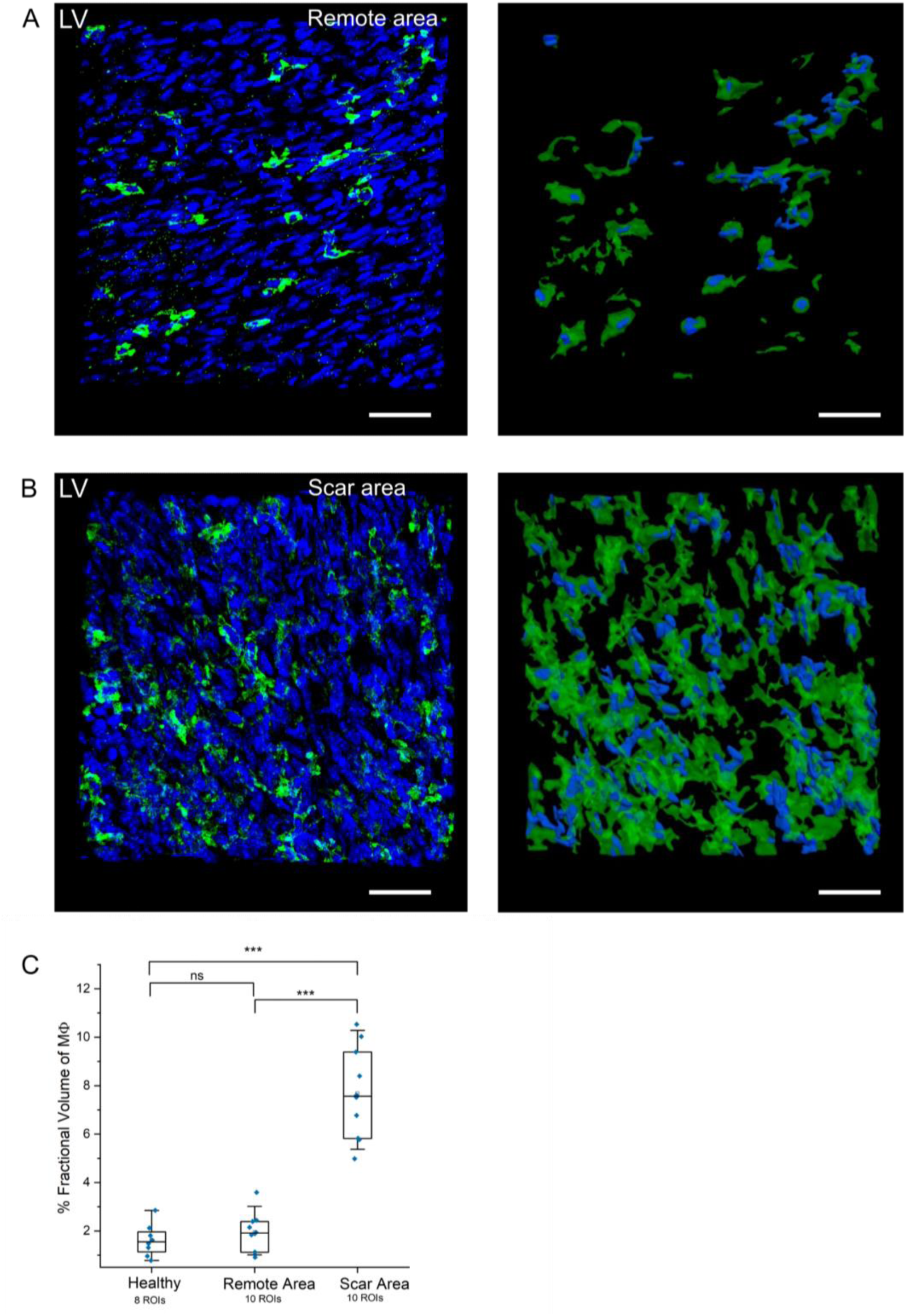
3D projections and reconstructions of MΦ in LV of cryo-ablated heart. 3D projection of confocal image stack (left) and 3D reconstruction (right) of ChR2-eYFP expressing MΦ in remote (A) and scar (B) regions, in a cryo-ablated Cx3cr1-ChR2 heart (scale bars – 50 μm). C) Fractional volume occupied by MΦ in LV regions of healthy hearts (n=8 ROI, N= 3 Cx3cr1-ChR2 hearts), compared to remote (n=10 ROI) and scar regions (n=10 ROI) of cryo-ablated hearts (N=3 Cx3cr1-ChR2 cryo-ablated hearts).

## 4. Discussion

While the relevance of cardiac NM for heart function in health and disease is being increasingly recognised (Lother & Kohl, 2023), there is still limited knowledge about the 3D structure of FB and MΦ in native myocardium. However, quantitative information on their distribution, morphology and dimension in healthy tissue is key to understanding how NM structure is changed in disease, an important prerequisite for developing NM-targeted interventions. In this study, we developed a combined wet and dry-lab approach to quantitatively assess the 3D structure of genetically labelled NM, both in healthy and pathologically remodelled mouse hearts. This includes electrophoretic tissue clearing by an adapted X-CLARITY protocol, which was applied to all chambers of the isolated mouse heart, followed by antibody staining for amplifying fluorescent signals from reporter proteins (here only necessary for eYFP-labelled MΦ) and nuclei staining. After incubation in refractive index matching mounting medium, large tissue volumes were imaged at high resolution by confocal microscopy. Finally, a custom developed algorithm was used to generate 3D reconstructions of genetically labelled NM and dye-stained nuclei, which allowed us to assess NM distribution in tissue, and the morphology of single cells and/or cell networks across the different chambers of the heart. 3D reconstructions were further used to calculate surface area and volume of individual cells as well as the fractional tissue volume occupied by each cell type. Based on these technical developments, we were able to shine light on yet unresolved 3D properties of FB and MΦ, assessing structural differences between the atria and ventricles, as well as during cardiac scar formation following ventricular cryo-injury. Our key findings are: (1) Cardiac FB and MΦ are abundant in all chambers of the heart, but FB generally occupy larger tissue volumes. (2) In healthy myocardium, FB form extended, interconnected networks, whereas the majority of MΦ appears as solitary cells. (3) FB and MΦ not only have similar morphologies, but also same-order-of-magnitude dimensions across all chambers of the heart. (4) Cardiac scar tissue is characterised by strongly altered tissue structure, including dense clusters of both FB and MΦ. (5) In myocardium remote to ventricular scars, the volume occupied by FB is significantly increased, while no change was observed in MΦ fractional volume, when compared to healthy myocardium.

### Distribution and morphology of FB and MΦ across murine heart chambers

Comparing both large 2D overview images and 3D reconstructions, FB and MΦ were abundant throughout the heart, but showed different distributions depending on cell type and cardiac chamber. While FB form large interconnected networks (containing up to 27 nuclei in the ventricles and up to 73 nuclei in the atria), most reconstructed MΦ were mononuclear, thus not in direct contact with neighbouring MΦ. This is in line with the primary functions of the individual cell types: While FB maintain the structural and mechanical integrity of the myocardium via maintenance and alteration of the extracellular matrix (ECM) (Gourdie *et al*., 2016; Klesen *et al*., 2018; Plikus *et al*., 2021), MΦ use phagocytosis to clean-up apoptotic cells and extracellular debris, relying on the ability to scavenge the cardiac interstitial space (Epelman *et al*., 2014; Zaman & Epelman, 2022). FB and MΦ have been demonstrated to interact with various cardiac cell types using classical paracrine signalling as well as biophysical cross-talk. The latter includes heterocellular electrotonic coupling and mechanical interactions, either based on direct cell-cell contact or relying on integrin-ECM bridging (Perbellini *et al*., 2018; Simon-Chica *et al*., 2023*b*; Lother & Kohl, 2023). In the present study, we observed finger-like micro-protrusions extending from both FB and MΦ, resembling tunneling nanotubes identified with electron microscopy in cryo-injured murine hearts (Quinn *et al*., 2016), potentially providing means of homo- and heterotypic interaction.

The morphology and distribution of NM networks is different between the ventricles and the atria. In the ventricles, FB networks are regularly structured and characterised by straight and orthogonal connections between adjacent FB, likely reflecting the high degree of CM alignment. In the atria, however, FB networks have more diffuse structures, exhibiting a combination of tortuous and straight patterns. Of note, the complexity of network structures can only be fully appreciated based on high-resolution 3D reconstructions, allowing insights into anisotropic tissue structure and cell-cell contacts, often missing in 2D images of myocardial tissue sections. Similar to FB, MΦ distribution and alignment follows the overall myocardial architecture, with stair-like MΦ wrapping around regularly aligned CM in the ventricles, and more diffusely distributed cells in the atria. Notably, we also observed small networks of MΦ next to CM, consistent with the reported phagocytosis of dysfunctional mitochondria and other cargo contained in ‘exophers’ released from healthy CM, in a process referred to as ‘heterophagy’ (Nicolás-Ávila *et al*., 2020, 2022).

### Quantitative assessment of NM dimensions in the healthy murine heart

Cardiac computational modelling has greatly advanced our comprehension of structure-function relationships and biophysical cell-cell cross-talk (Quinn & Kohl, 2013), with the possibility of linking data obtained in preclinical animal studies to clinical observations in patients (Niederer *et al*., 2019). Specifically, mechanistic modelling has promoted our understanding of heterocellular electrical coupling, and its relevance for whole-organ electrophysiology (Kohl *et al*., 2005; Hulsmans *et al*., 2017; Simon-Chica *et al*., 2022, 2023*b*; Wang *et al*., 2023). Parametrisation of *in-silico* models of cardiac electrophysiology requires quantitative experimental data as foundation, including passive and active electrical properties of the considered cell types. Accordingly, the dimensions of FB and MΦ in tissue are important, as the membrane surface area is directly proportional to the electrical capacitance and inversely correlated with membrane resistance of NM (Simon-Chica *et al*., 2022), and both are essential parameters to computationally predict effects of NM-CM coupling. Classically, these parameters have been measured by electrophysiological recordings of isolated cells, with large discrepancies to cell behavior *in situ*. The surface area and capacitance of freshly isolated FB have previously been determined to be 150–250 μm^2^ and 4.5-6 pF (Dawson *et al*., 2012; Kohl & Gourdie, 2014). In contrast, using electron microscopy, the FB surface area was 720 μm^2^ *in situ* (De Mazière *et al*., 1992), which still may underestimate actual dimensions as the study was based on 2D images and did not include distal FB extensions. By comparison, the average surface area of FB in healthy LV was 1560±130 µm^2^ in the current study. A similar discrepancy is found for MΦ. Reported values for surface area and capacity of freshly isolated MΦ are 380 μm^2^ and 18 pF (Ongstad & Kohl, 2016). Hulsmans *et al*. reported a surface area of 149±24 µm^2^ for MΦ in the atrioventricular node *in situ* in hearts from Cx3cr1-ChR2 mice, however; this value was later identified to be the area of a maximum intensity projection, rather than corresponding to the actual membrane surface (Hulsmans *et al*., 2017). We previously reported an average MΦ surface area of 1160±80 μm^2^ (Simon-Chica *et al*., 2022), whereas here we determined a surface area of 1460±100 μm^2^, both in healthy murine LV tissue. The discrepancy between the two studies can be explained by the used FP. While the earlier study used a FP present in the cytoplasm, we here used ChR2-eYFP, leading to efficient labelling of the plasmalemma. This allows detection of fine membrane protrusions not visible with the cytoplasmic marker, thus resulting in larger values of surface area. Our study thus provides important quantitative data to estimate the passive electrophysiological properties of FB and MΦ *in-situ*, paving the way to establishing more realistic *in-silico* models of heterocellular biophysical interactions.

### Structural alterations in ventricular scar tissue

FB and MΦ are key players involved in tissue remodelling and scarring in injured hearts, such as following myocardial infarction. Early during cardiac scar formation, FB are activated and trans-differentiate into myofibroblasts (myoFB), characterised by a migratory phenotype and expression of contractile microfilaments (Rog-Zielinska *et al*., 2016), and with diverse functions in tissue repair, including regulation of ECM dynamics, vascularisation and immune responses (Plikus *et al*., 2021). Following myocardial injury, the MΦ population is expanded by both local proliferation and monocyte recruitment (Heidt *et al*., 2014), and scar-resident MΦ orchestrate inflammation and mediate adaptive remodelling to changes in tissue mechanics (Wong *et al*., 2021). Furthermore MΦ are, in part, responsible for activation of FB via secretion of pro-inflammatory signalling molecules, such TGF-β, and directly contribute to deposition of scar-forming collagen (Simões *et al*., 2020). In the current study, we quantitatively assessed changes in the 3D distribution and structure of FB and MΦ in scar tissue 28 days after ventricular cryoinjury (Jensen *et al*., 1987). We observe that both FB and MΦ numbers are increased in ventricular scar tissue, indicative of NM recruitment and local proliferation. Accordingly, the FV occupied by FB and MΦ are 4.9-fold and 3-fold higher in the scar area, respectively. Surprisingly, the FV of FB is 1.7-fold increased also in ventricular areas remote to the scar. A longitudinal observation could reveal whether remote FB expansion is occurring during the acute inflammatory phase or rather during later stages of scar formation, *i.e.* as a secondary effect to altered mechanical load caused by the compromised ventricular contractility. FV occupied by MΦ, on the contrary, was not different between remote and healthy LV.

Whereas FB networks in tissue regions remote to scars showed similar network structures as in healthy myocardium, FB networks within scars were dense, irregularly structured and contained many interconnecting cell extensions. In the scar area, MΦ also form extended networks, such that individual MΦ cannot be separated from one another using our segmentation algorithm. Ventricular cryo-injury is characterised by temperature-induced necrosis of all cell types (Jensen *et al*., 1987), leading to loss of tissue structuring alignment of cardiomyocytes, vessels and nerve fibers. In addition, during the acute inflammatory phase, interstitial ECM is degraded, followed by the production and resolution of the provisional ECM (proliferative phase), and, finally, mature scar formation (maturation phase) (Silva *et al*., 2021). Accordingly, scar-forming NM replace cell debris and ECM forming diffuse, interconnected cell networks lacking the characteristic structural alignment of healthy ventricles, with direct consequences on scar mechanics (Fomovsky & Holmes, 2010) and trans-scar electrical conduction (Rog-Zielinska *et al*., 2016).

### Experimental challenges and future perspectives

The here presented experimental approach is based on cell-type specific genetic labelling, tissue clearing, confocal imaging of large tissue volumes, and 3D segmentation of nuclei-containing cell networks. Each of these steps comes with its intrinsic challenges and/or limitations. Tamoxifen-inducible Cre recombination mouse models offer unprecedented means for fluorescent labelling of cell populations at defined time points in respect to cardiac events, but rely on the availability of highly specific and efficient recombination in the target cells (Feil *et al*., 1996; Acharya *et al*., 2011; Madisen *et al*., 2012; Yona *et al*., 2013). Genetic labelling can be combined with immunofluorescence staining, but established protocols for antibody labelling of cleared myocardium are time-consuming, thereby hampering further optimisation (Kolesová *et al*., 2021). Similarly, while several methods for optical clearing of myocardial tissue and whole-hearts have been tested and further adjusted, their application remains challenging, especially in terms of preservation of tissue integrity and FP while aiming to achieve transmural transparency (Kolesová *et al*., 2021; Fischesser *et al*., 2021; Olianti *et al*., 2022). In the current study, we performed high-resolution confocal microscopy for imaging membrane-bound eYFP fluorescence, however; this approach is unsuitable for imaging whole murine hearts, which might be plausible using optimised light-sheet microscopes such as the mesoSPIM platform, enabling fast volumetric imaging of centimetre-sized cleared samples with near-isotropic resolution (Voigt *et al*., 2019). Finally, we developed a semi-automated algorithm to obtain 3D segmentations of eYFP and nuclei-labelled NM. A key issue with the algorithm is the remaining need for user intervention due to non-homogenous brightness between samples. Furthermore, assessing colocalisation can be challenging due to limited overlap between fluorescent signals from labelled nuclei and plasma membranes. The use of cytoplasmic indicators would ease this, but would prevent tracking small membrane protrusions. The prevalence of extended FB networks meant that quantification of FB surface area and volume was done by dividing network size by the number of nuclei found in the network, rather than direct measurements of single FB, thus providing average values per network rather than insights into the size distribution and variance between individual cells. Similarly, identification of single MΦ was complex in the atria and extraordinarily hard in the scar of the cryo-ablated hearts, where segmentation of individual nuclei was challenging by itself. Previous work has used the rainbow reporter system crossed with tamoxifen-dependent Cre lines, to follow clonal expansion in the developing mouse heart, which helped to dissect CM proliferative capacity and dynamics during embryonic and post-natal development (Sereti *et al*., 2018). Combining the rainbow system for multi-colour labelling of cardiac NM with optimised tissue clearing, may be used for future assessment of cell-cell contacts between individual NM in complex cell networks, as found in healthy atria and in ventricular scar tissue. Our results indicate that membrane-bound eYFP is especially well suited for revealing small membranous protrusions of NM. The exact nature and role of these protrusions is uncertain, but they are likely to play a role in intercellular communication and/or sensing of the surrounding environment.

### Concluding statement

Our study provides an atlas of the distribution, morphology and quantitative dimensions of FB and MΦ across all chambers of the healthy murine heart, as well as in scarred ventricles following left-ventricular cryo-injury. This may serve as a basis for future investigations relating NM structure and function, and assessing biophysical cell-cell interactions, both crucial aspects for defining NM contributions to cardiac physiology in health and disease.

## Acknowledgments

We thank the technical support team of IEKM for excellent assistance. We thank all members of IEKM for critical discussion of the research and manuscript. We acknowledge the microscopy facility SCI-MED (Super-Resolution Confocal/ Multiphoton Imaging for Multiparametric Experimental Designs) at IEKM, Freiburg, for providing access to imaging setups and analysis work stations.

This work was funded by the German Research Foundation (DFG; #412853334 to FSW). All authors are members of SFB1425, funded by the DFG (#422681845), and FSW is associated with the local Cluster of Excellence CIBSS (#390939984).

