## Supplementary Material for "3D structure of fibroblasts and macrophages in the healthy and cryo-ablated heart"

**Supplementary Data**

**Supplementary Figure 1.** Graphical abstract summarising mouse models used to assess FB and MΦ structure in healthy and cryo-ablated hearts.

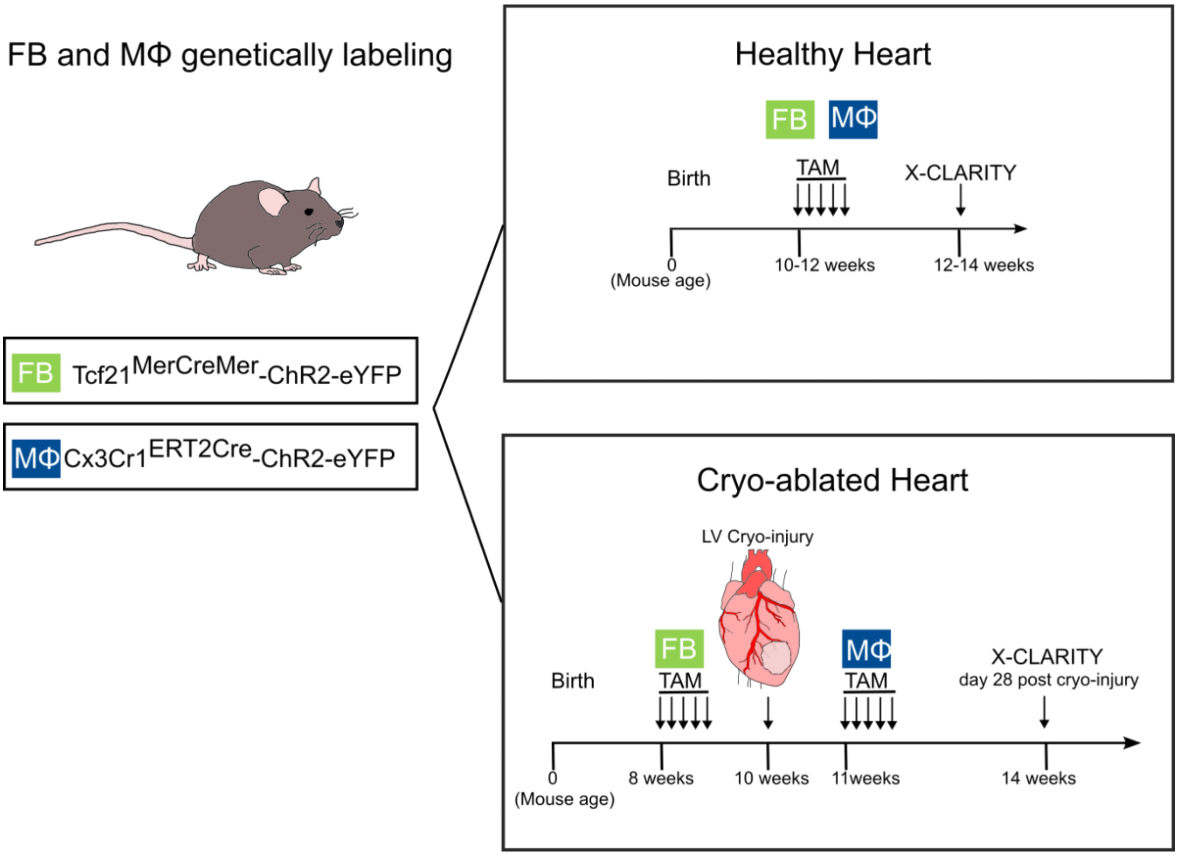


**Supplementary video 1**. Z-track of confocal microscopy images and corresponding 3D reconstruction of FB networks in ROI of cleared healthy LV.

**Supplementary video 2.** Z-track of confocal microscopy images and corresponding 3D reconstruction of MΦ in ROI of cleared healthy LV.

**Supplementary video 3.** Z-track of confocal microscopy images and corresponding 3D reconstruction of FB networks in the scar of cleared cryo-ablated LV.

**Supplementary video 4.** Z-track of confocal microscopy images and corresponding 3D reconstruction of MΦ networks in the scar of cleared cryo-ablated LV.

**Supplementary tables**

**Table1**. Values of voxel size of 3D reconstructions of FB and MΦ from LV, RV, RA, LA and septum in cleared healthy hearts.

| **Chamber** | **nROIs** | **FB** | | **MΦ** | |
|---|---|---|---|---|---|
| | | X,Y (μm) | Z( μm) | X,Y (μm) | Z(μm) |
| **LV** | 1 | 0.05 | 0.37 | 0.13 | 0.29 |
| **LV** | 2 | 0.05 | 0.38 | 0.13 | 0.29 |
| **LV** | 3 | 0.05 | 0.39 | 0.13 | 0.24 |
| **LV** | 4 | 0.04 | 0.31 | 0.13 | 0.29 |
| **LV** | 5 | 0.04 | 0.28 | 0.12 | 0.29 |
| **LV** | 6 | 0.04 | 0.28 | 0.11 | 0.29 |
| **LV** | 7 | 0.04 | 0.91 | 0.11 | 0.29 |
| **LV** | 8 | 0.05 | 0.91 | 0.11 | 0.29 |
| **LV** | 9 | 0.05 | 0.37 | | |
| **LV** | 10 | 0.04 | 0.37 | | |
| **LV** | 11 | 0.11 | 0.29 | | |
| **RV** | 12 | 0.04 | 0.33 | 0.12 | 0.29 |
| **RV** | 13 | 0.04 | 0.36 | 0.13 | 0.29 |
| **RV** | 14 | 0.06 | 0.33 | 0.27 | 0.24 |
| **RV** | 15 | 0.06 | 0.33 | 0.13 | 0.24 |
| **RV** | 16 | 0.06 | 0.33 | 0.13 | 0.29 |
| **RV** | 17 | 0.04 | 0.31 | 0.13 | 0.29 |
| **RV** | 18 | 0.04 | 0.38 | | |
| **RV** | 19 | 0.05 | 0.38 | | |
| **RV** | 20 | 0.04 | 0.36 | | |
| **RV** | 21 | 0.05 | 0.38 | | |
| **RV** | 22 | 0.11 | 0.29 | | |
| **RV** | 23 | 0.11 | 0.29 | | |
| **RV** | 24 | 0.11 | 0.29 | | |
| **LA** | 25 | 0.04 | 0.38 | 0.95 | 0.30 |
| **LA** | 26 | 0.04 | 0.38 | 0.95 | 0.30 |
| **LA** | 27 | 0.04 | 0.38 | 0.13 | 0.34 |
| **LA** | 28 | 0.04 | 0.38 | 0.13 | 0.29 |
| **LA** | 29 | 0.04 | 0.38 | 0.13 | 0.34 |
| **LA** | 30 | 0.04 | 0.37 | 0.13 | 0.34 |
| **LA** | 31 | 0.11 | 0.29 | | |
| **LA** | 32 | 0.11 | 0.29 | | |
| **LA** | 33 | 0.11 | 0.29 | | |
| **RA** | 34 | 0.05 | 0.38 | 0.13 | 0.29 |
| **RA** | 35 | 0.05 | 0.38 | 0.13 | 0.30 |
| **RA** | 36 | 0.04 | 0.39 | 0.13 | 0.29 |
| **RA** | 37 | 0.04 | 0.37 | 0.13 | 0.29 |
| **RA** | 38 | 0.06 | 0.27 | 0.13 | 0.36 |
| **RA** | 39 | 0.06 | 0.27 | | |
| **RA** | 40 | 0.11 | 0.29 | | |
| **RA** | 41 | 0.11 | 0.29 | | |
| **RA** | 42 | 0.11 | 0.29 | | |
| **Septum** | 43 | 0.04 | 0.38 | 0.13 | 0.29 |

| | | | | | |
|---|---|---|---|---|---|
| **Septum** | 44 | 0.05 | 0.33 | 0.13 | 0.29 |
| **Septum** | 45 | 0.07 | 0.35 | 0.13 | 0.29 |
| **Septum** | 46 | 0.05 | 0.39 | 0.13 | 0.29 |
| **Septum** | 47 | 0.07 | 0.35 | | |
| **Septum** | 48 | 0.11 | 0.29 | | |
| **Septum** | 49 | 0.11 | 0.29 | | |
| **Septum** | 50 | 0.11 | 0.29 | | |
| **Septum** | 51 | 0.11 | 0.29 | | |

**Table 2**. Values of voxel size of 3D reconstructions of FB and MΦ from LV in cryo-ablated hearts.

| **ROIs** | **FB** | | **MΦ** | |
|---|---|---|---|---|
| | X,Y (μm) | Z(μm) | X,Y(μm) | Z(μm) |
| **Remote-LV** | 0.07 | 0.28 | 0.14 | 0.29 |
| **Remote-LV** | 0.14 | 0.29 | 0.14 | 0.29 |
| **Remote-LV** | 0.07 | 0.28 | 0.14 | 0.29 |
| **Remote-LV** | 0.07 | 0.29 | 0.14 | 0.29 |
| **Remote-LV** | 0.14 | 0.28 | 0.14 | 0.29 |
| **Remote-LV** | 0.14 | 0.28 | 0.14 | 0.28 |
| **Remote-LV** | 0.14 | 0.28 | 0.14 | 0.28 |
| **Remote-LV** | 0.14 | 0.28 | | |
| **Remote-LV** | 0.14 | 0.28 | | |
| **Remote-RV** | 0.14 | 0.28 | 0.14 | 0.28 |
| **Remote-RV** | 0.14 | 0.28 | 0.14 | 0.28 |
| **Remote-RV** | 0.14 | 0.28 | 0.14 | 0.28 |
| **Remote-RV** | 0.14 | 0.28 | | |
| **Remote-RV** | 0.14 | 0.28 | | |
| **Remote-RV** | 0.14 | 0.28 | | |
| **Scar-LV** | 0.07 | 0.29 | 0.14 | 0.29 |
| **Scar-LV** | 0.07 | 0.28 | 0.14 | 0.29 |
| **Scar-LV** | 0.07 | 0.28 | 0.14 | 0.29 |
| **Scar-LV** | 0.07 | 0.42 | 0.14 | 0.29 |
| **Scar-LV** | 0.07 | 0.29 | 0.14 | 0.29 |
| **Scar-LV** | 0.07 | 0.28 | 0.14 | 0.28 |
| **Scar-LV** | 0.07 | 0.28 | 0.14 | 0.28 |
| **Scar-LV** | 0.11 | 0.29 | 0.14 | 0.28 |
| **Scar-LV** | 0.14 | 0.28 | 0.14 | 0.28 |
| **Scar-LV** | 0.14 | 0.28 | 0.14 | 0.28 |
| **Scar-LV** | 0.14 | 0.28 | | |
| **Scar-LV** | 0.14 | 0.28 | | |
| **Scar-LV** | 0.14 | 0.28 | | |
| **Scar-LV** | 0.14 | 0.28 | | |

Table 3. Values of area per cell type in respect to the total tissue area, obtained from 2D reconstructions of FB and MΦ in Tcf21- and Cx3cr1-ChR2-eYFP hearts, respectively.

| **Chambers** | **FB** | | **MΦ** | | |
|---|---|---|---|---|---|
| | Tissue Total Area ($x10^6$ $\mu m^2$) | Cell Total Area ( $x10^5 \mu m^2$) | Tissue Total Area ( $x10^6$ $\mu m^2$) | Cell Total Area ( $x10^4$ $\mu m^2$) | # Cells in Tissue |
| **LV** | 8.3 | 3.4 | 7.7 | 16.5 | 1276 |
| **LA** | 3.4 | 8.4 | 6.3 | 9.5 | 606 |
| **RA** | 4.6 | 4.7 | 5.9 | 8.9 | 1374 |
